# Self-organization mechanism of distinct microtubule orientations in axons and dendrites

**DOI:** 10.1101/163014

**Authors:** Honda Naoki, Katsuhiro Uegaki, Shin Ishii

## Abstract

The identities of axons and dendrites are acquired through the self-organization of distinct microtubule (MT) orientations during neuronal polarization. The axon is generally characterized by a uniform MT orientation with all plus-ends pointing outward to the neurite terminal (‘plus-end-out’ pattern). On the other hand, the MT orientation pattern in the dendrites depends on species: vertebrate dendrites have a mixed alignment with both plus and minus ends facing either the terminal or the cell body (‘mixed’ pattern), whereas invertebrate dendrites have a ‘minus-end-out’ pattern. However, how MT organizations are developed in the axon and the dendrites is largely unknown. To investigate the mechanism of MT organization, we developed a biophysical model of MT kinetics, consisting of polymerization/depolymerization and MT catastrophe coupled with neurite outgrowth. The model simulation showed that the MT orientation can be controlled mainly by the speed of neurite growth and the hydrolysis rate. With a low hydrolysis rate, vertebrate plus-end-out and mixed microtubule patterns emerged in fast- and slow-growing neurites, respectively. In contrast, with a high hydrolysis rate, invertebrate plus-end-out and minus-end-out microtubule patterns emerged in fast- and slow-growing neurites, respectively. Thus, our model can provide a unified understanding of distinct microtubule organizations by simply changing the parameters.

## Background

Neurons are highly polarized cells consisting of functionally distinct compartments, the axon and the dendrites. During development, neurons initially extend multiple immature neurites, which undergo repeated protrusion and retraction but are on average symmetric in length. Then, the symmetry suddenly breaks with the rapid growth of a randomly selected neurite [1] (**Fig. 1A**). The growing neurite (major neurite) becomes an axon, which subsequently migrates to connect with the target neurons [2], whereas the remaining neurites (minor neurites) grow slowly, thereby developing into dendrites. However, how the major and minor neurites, which differ only in length and growth speed, acquire the different identities of the axon and the dendrites remains elusive.

**Figure 1:**
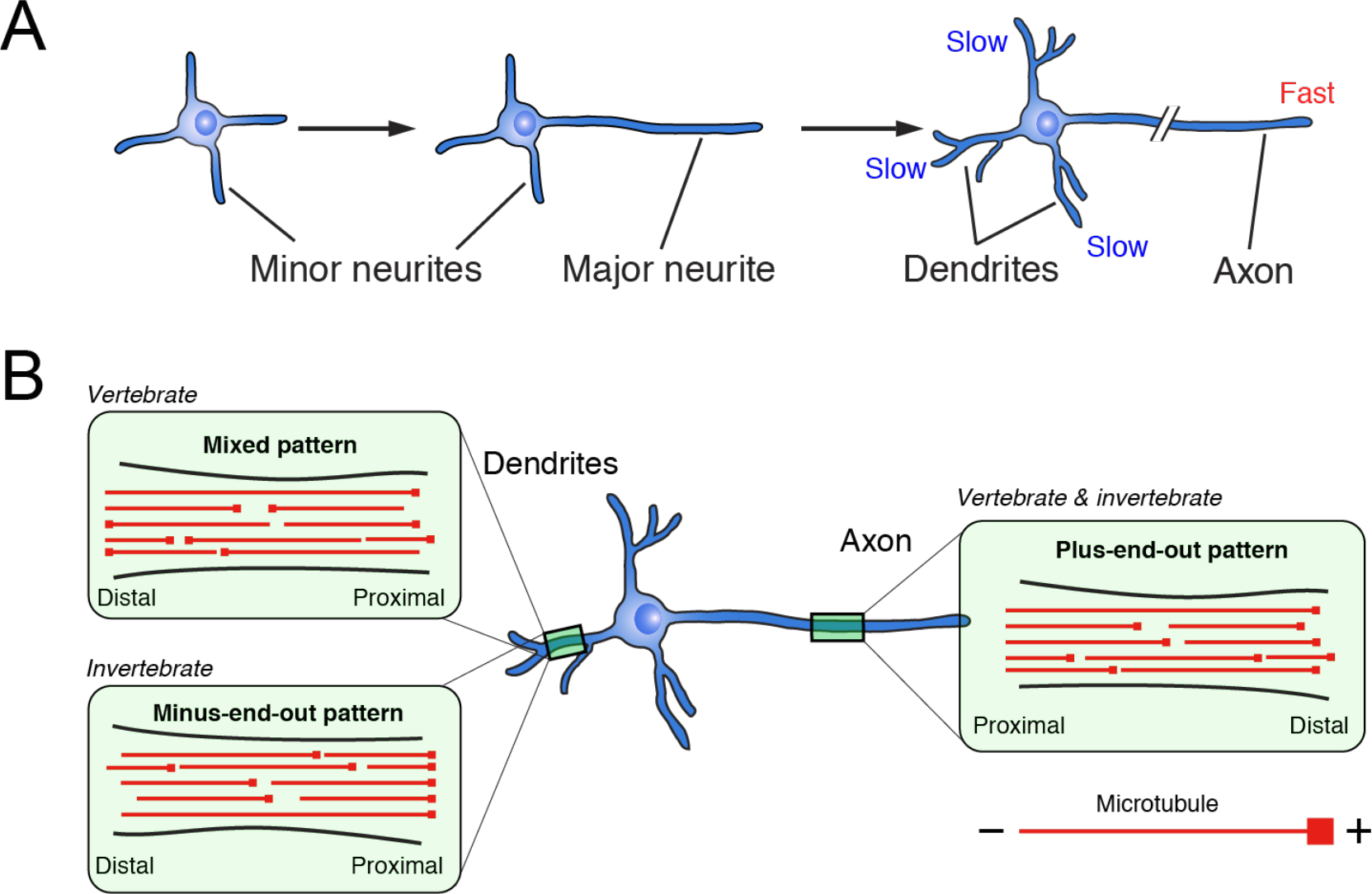
Self-organization of MT orientations acquired through neuronal polarization. **(A)** Developmental process of neuronal polarization. Neurons initially extend several immature neurites, which are symmetric in length and are not committed to axon or dendrites. One neurite then breaks the initial morphological symmetry with rapid growth. The growing neurite (major neurite) becomes an axon, whereas the remaining ones (minor neurites) slowly develop into dendrites. **(B)** MT orientations in axon and dendrites. In vertebrates, the axon and dendrites contain plus-end-out and mixed MT orientations, respectively. In invertebrates, the axon and dendrites contain plus-end-out and minus-end-out MT orientations, respectively.

The identities of the axon and dendrites are characterized by structural differences in microtubule (MT) orientation [3,4] (**Fig. 1B**), which affect the direction of MT-based motor proteins [5,6]. Many studies have investigated the MT orientations in the axon and dendrites using electron microscopy [7], second-harmonic generation microscopy [8], or the live imaging of fluorescently labeled MT plus-end-binding proteins [9,10]. In various types of vertebrate neurons, the axon has a uniform MT orientation with all plus ends pointing outward to the terminal (‘plus-end-out’ pattern), whereas dendrites have a mixed orientation with both plus and minus ends facing either the terminal or the cell body (‘mixed’ pattern). In invertebrate neurons, axons also exhibit a plus-end-out MT orientation. Interestingly, in contrast, invertebrate dendrites have a uniform MT orientation with all minus ends pointing outward to the terminal (‘minus-end-out’ pattern) [11,12]. Upon neuronal polarization, the MT orientation in immature neurites starts with a mixed pattern in which the fraction of plus-end-out MTs is dominant and then gradually self-organizes to a plus-end-out pattern in the axon and a fifty-fifty mixed pattern in the dendrites [10], suggesting that the first crucial step for the acquisition of axon and dendrite identities is distinct MT orientations. However, the self-organization mechanism of the three distinct MT orientation patterns (i.e., plus-end-out, minus-end-out, mixed) in the axon and dendrites is largely unclear.

Neuronal polarization has been extensively investigated with computational models [13–15], including ours [16,17], but all previous models focused on morphological symmetry breaking. While these models provided insights into the mechanism by which only a single neurite among immature minor neurites is selected to grow, none addressed how the growing major neurite and remaining minor neurites acquire the identities of the axon and dendrites.

In this study, we sought to determine the MT orientation-based mechanism underlying the acquisition of the axon and dendrite identities. By developing a biophysical computational model of MT kinetics in developing neurites, we investigated how the self-organization of MT orientation is affected by the balance among the polymerization/depolymerization rates of MT plus and minus ends, the hydrolysis rate and the growth rate of the neurites. This model demonstrated that different MT orientations emerged depending on the parameters of the MT kinetics, which presents a unified view of the plus-end-out, minus-end-out and mixed pattern formations.

## Results

### Model of MT kinetics in growing neurites

To examine the self-organization mechanism of distinct MT orientations in the axon and dendrites, we developed a biophysical model of the MT assembly in a growing neurite. In the model, MTs align one-dimensionally along the neurite shaft (**Fig. 2A**). The MTs independently elongate and shrink in the growth cone located at the neurite tip, where the MT elongation is bounded by the distal end of the neurite. The model neurite grows at a speed of *v_n_*, which makes additional space for MT elongation. For the sake of simplicity, *v_n_* is a control parameter independent of the MTs, as the neurite growth is predominantly driven by actin filament (F-actin) [18,19], although the MTs interact with and support F-actin in the growth cone [20]. Note that major and minor neurites grow rapidly and slowly, respectively, during neuronal polarization. Since MT fragments are nucleated in the cell body and actively transported to the growth cones [3], existing MTs are replaced with MT fragments in the growth cone, where their orientations are selected at random. This MT replacement occurs in two situations: 1. any time, spontaneously, at a rate of *k_rep_* and 2. when the MT shrinks and passes outside the growth cone to the neurite shaft. Because the MTs are linked to each other by MT-associated proteins and stabilized as MT bundles [3], we did not consider MT elongation toward the cell body.

**Figure 2:**
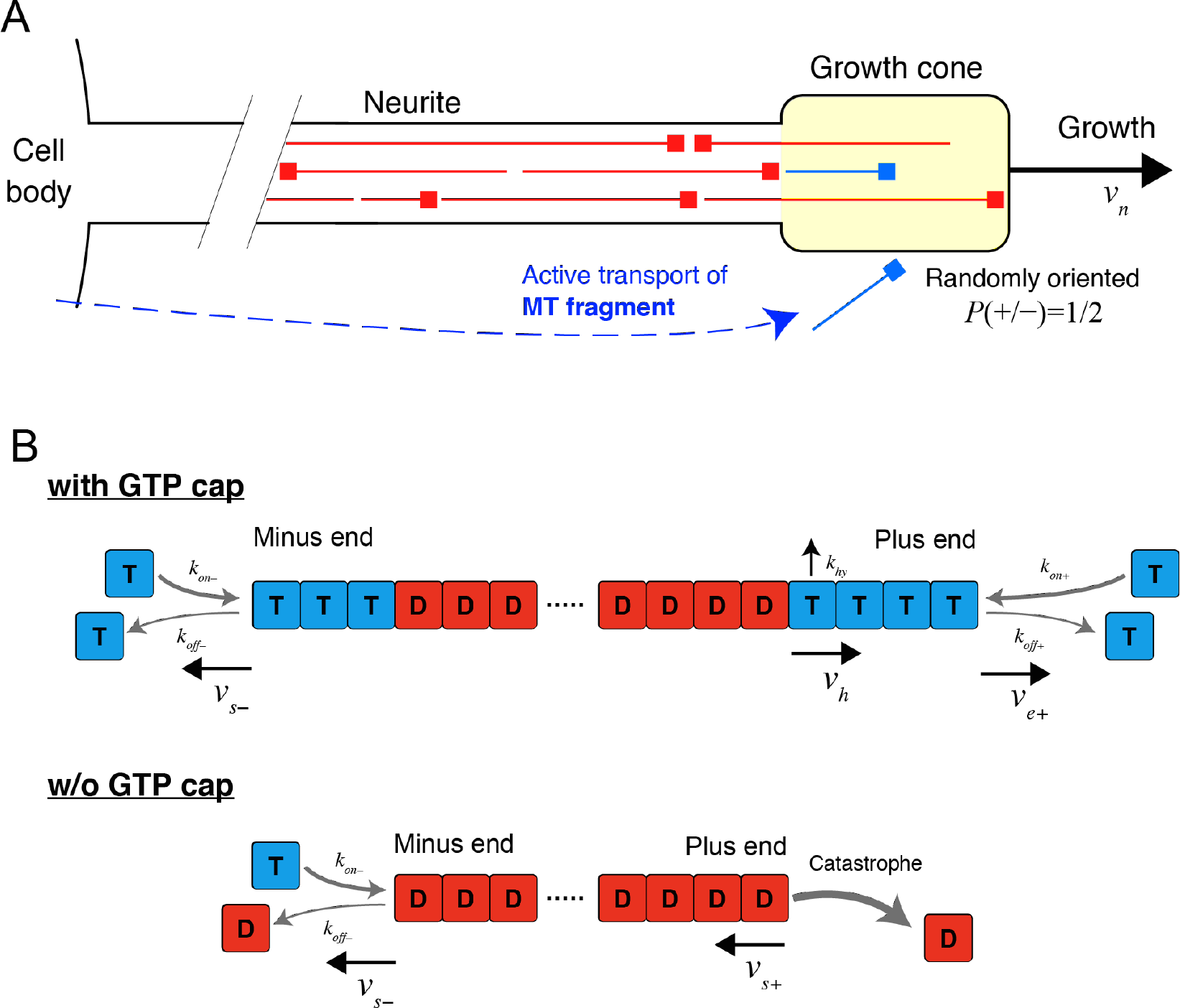
Biophysical model of MT kinetics in a growing neurite. **(A)** Schematic of MT assembly dynamics. MTs align along the neurite and elongate and shrink in the growth cone. The neurite grows at a speed of *v_n_*. The MTs are replaced with actively transported MT fragments (blue segment) in the growth cone, stochastically at rate of *k_rep_* and deterministically when the MT shrinks to lie outside the growth cone. **(B)** Schematic of MT reaction kinetics. The MTs polymerize GTP-bound tubulins and depolymerize at both ends, where the GTP-bound tubulins are converted to the GDP-bound form by hydrolysis. The loss of the GTP cap at the plus end initiates catastrophic shrinkage of the MT, whereas the minus end is stable in the absence of the GTP cap.

The MT elongates and shrinks following its reaction kinetics (**Fig. 2B**). The MT polymerizes and depolymerizes at both ends, where those rates are biased such that the MT elongates faster at the plus end than at the minus end. Tubulin heterodimers as monomers are added to the MT in the GTP-bound form and hydrolyze to the GDP-bound form at rate of *k_hy_*, and thus polymerizing MTs have a cap consisting of GTP-tubulin (GTP cap). Upon the loss of the GTP cap at the plus end, the MT suddenly undergoes quick depolymerization, called catastrophe [21]. The minus end does not undergo catastrophe, because the minus end is stabilized by minus-end-binding proteins such as CAMSAPs [22]. Although the MT is a tubular polymer consisting of 13 protofilaments [23], we addressed its kinetics above as a simple single filament following previous computational models [24–26] (see details in **Materials and Methods**).

### Dynamic properties of MT

We first investigated the dynamic properties of a single MT by simulation (**Fig. 3**). Suppose that the MT fragment was actively transported to the halted growth cone and incorporated into the pre-existing MT bundle with a random orientation. In this situation, the plus-end-out MT fragment rapidly elongated and reached the distal end of the neurite (**Fig. 3A**). Then, the GTP cap was shortened and removed via hydrolysis, which led to catastrophic shrinkage of the MT. The MT then disappeared from the growth cone. This transition from elongation to pause to shrinkage of the MT is consistent with the MT behaviors observed in the cell periphery [27,28]. The minus-end-out MT fragment also elongated to reach the distal end of the neurite, but this process was relatively slow compared with that of the plus end (**Fig. 3B**). Because the minus end of the MT is stabilized in vivo independently of the loss of the GTP cap [22], the minus-end-out MT did not show catastrophe and continued to exist.

**Figure 3:**
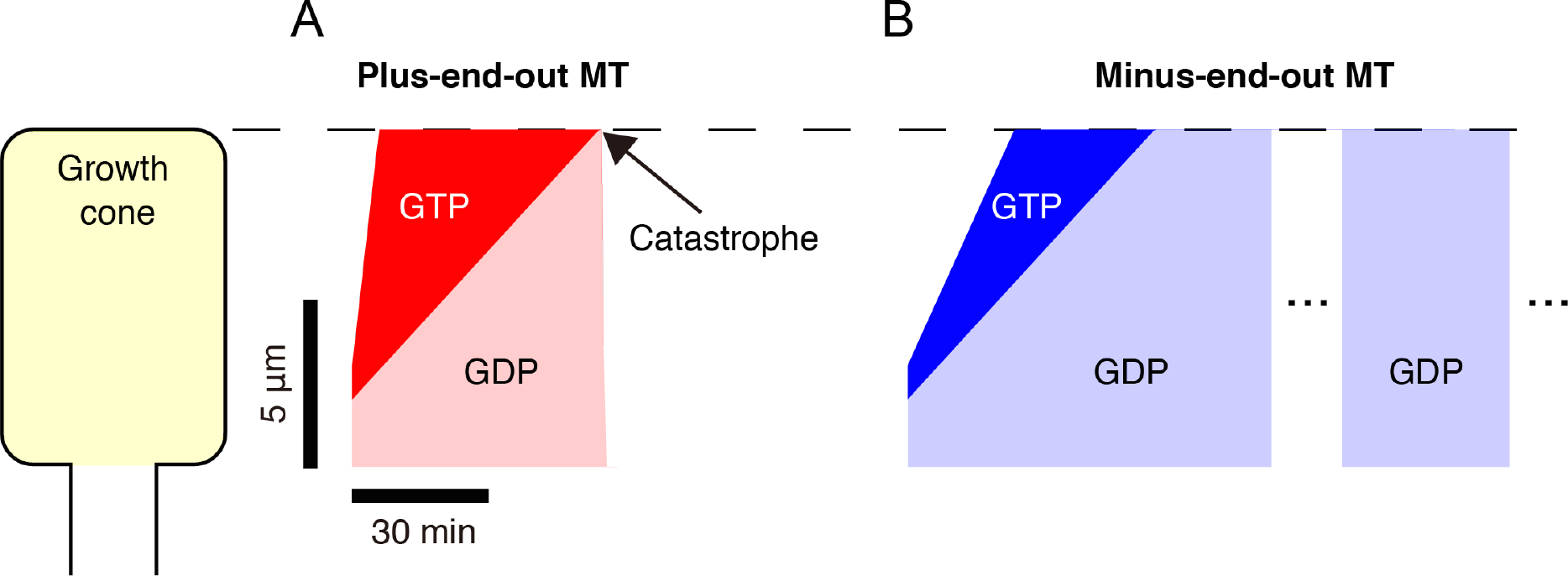
Dynamic properties of MT within a paused neurite. The temporal dynamics of a single plus-end-out MT **(A)** and minus-end-out MT **(B)** in the immobile neurite were simulated. The MT was initially located beneath the neurite terminal. The black dashed line indicates the position of the neurite terminal. The red and blue regions represent the GTP caps at the plus and minus ends, respectively, whereas the transparent regions represent the GDP-bound tubulins.

### MT orientation patterns in vertebrates

Next, we explored how the MT orientation pattern evolves in neurites growing at different speeds, as the major and minor neurites grow rapidly and slowly, respectively. In a fast-growing neurite, the plus-end-out MT caught up to the moving distal tip but did not undergo catastrophe because the speed of the GTP cap disappearance (hydrolysis speed) was slower than the speed of neurite growth (**Fig. 4A**). The plus-end-out MT was stochastically replaced with a randomly oriented MT fragment. The newly incorporated minus-end-out MT could not catch up to the moving distal tip and was excluded from the growth cone to remain in the neurite shaft, because of the low polymerization rate at the minus end compared with the plus end. To see the temporal evolution of the MT orientation, we simulated the MT population, beginning with a mixed pattern with a bias toward plus-end-out MTs, as seen in immature neurites [10]. We then observed that the fraction consisting of the plus-end-out MTs gradually became dominant in a fast-growing neurite, which corresponds to the plus-end-out pattern observed in the axon (**Fig. 4B, C**). In a slow-growing neurite, both plus-end-out and minus-end-out MTs caught up to the moving distal tip and were frequently replaced with randomly oriented MT fragments (**Fig. 4D**). In the MT population, we observed equal fractions of plus- and minus-end-out MTs, which corresponds to the mixed pattern observed in vertebrate dendrites (**Fig. 4E, F**). Therefore, these results suggested that the MT-based structural identification of the vertebrate axon and dendrites is determined by differences in the growth speed of major and minor neurites.

**Figure 4:**
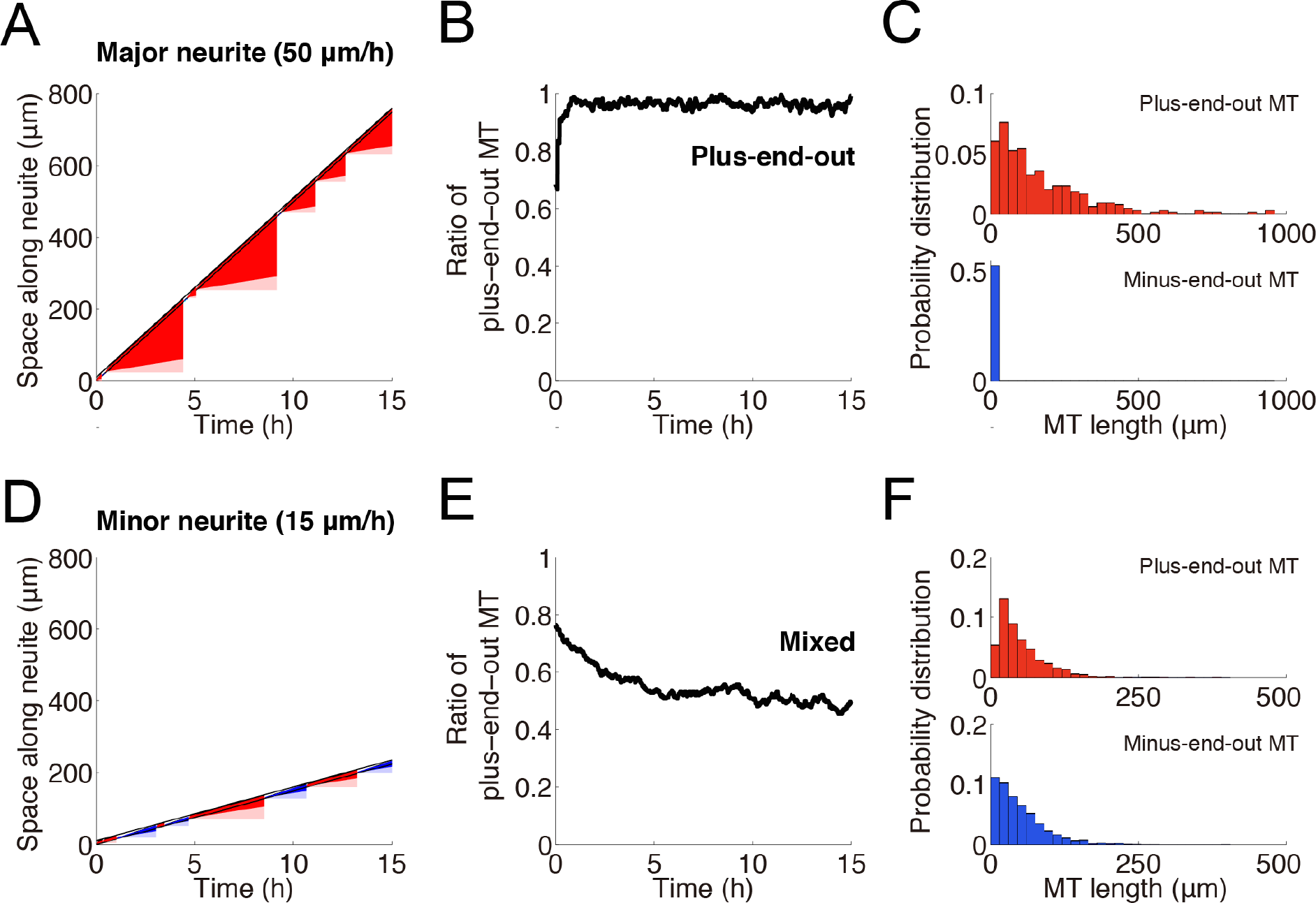
Self-organized MT orientations in vertebrate axon and dendrites. The temporal dynamics of MT replacement in fast- **(A-C)** and slow- **(D-F)** growing neurites were simulated with a low hydrolysis speed (8.86 (µm/h)). **(A, D)** Two black lines indicate the positions of the distal and proximal growth cone. Red and blue regions represent the GTP cap of the plus-end-out and minus-end-out MTs, respectively, whereas their transparent regions represent the GDP-bound tubulins. **(B, E)** The MT populations (n = 250) were simulated. The initial proportion of plus-end-out MTs was 75%. The black line indicates the ratio of the total length of plus-end-out MTs to the total length of all MTs. **(C, F)** The red and blue histograms represent the probability distributions of the length of the plus-end-out and minus-end-out MTs, respectively.

### MT orientation patterns in invertebrates

To further seek the mechanism of the minus-end-out pattern observed in invertebrate dendrites [11,12], we investigated the effect of hydrolysis on the self-organization of MT orientations. Increasing the hydrolysis rate in the model, we performed the same analysis as in Fig. 4. In a fast-growing neurite, we again obtained the plus-end-out MT orientation pattern observed in the axon (**Fig. 5A-C**). On the other hand, in a slow-growing neurite, both plus-end-out and minus-end-out MTs caught up to the moving distal tip, but only plus-end-out MTs were eliminated because hydrolysis was a faster process than the neurite growth (**Fig. 5D**). Thus, in the MT population, the fraction of minus-end-out MTs became dominant (**Fig. 5E, F**). These results indicated an important role of hydrolysis in the self-organization of the MT orientation patterns in invertebrates.

**Figure 5:**
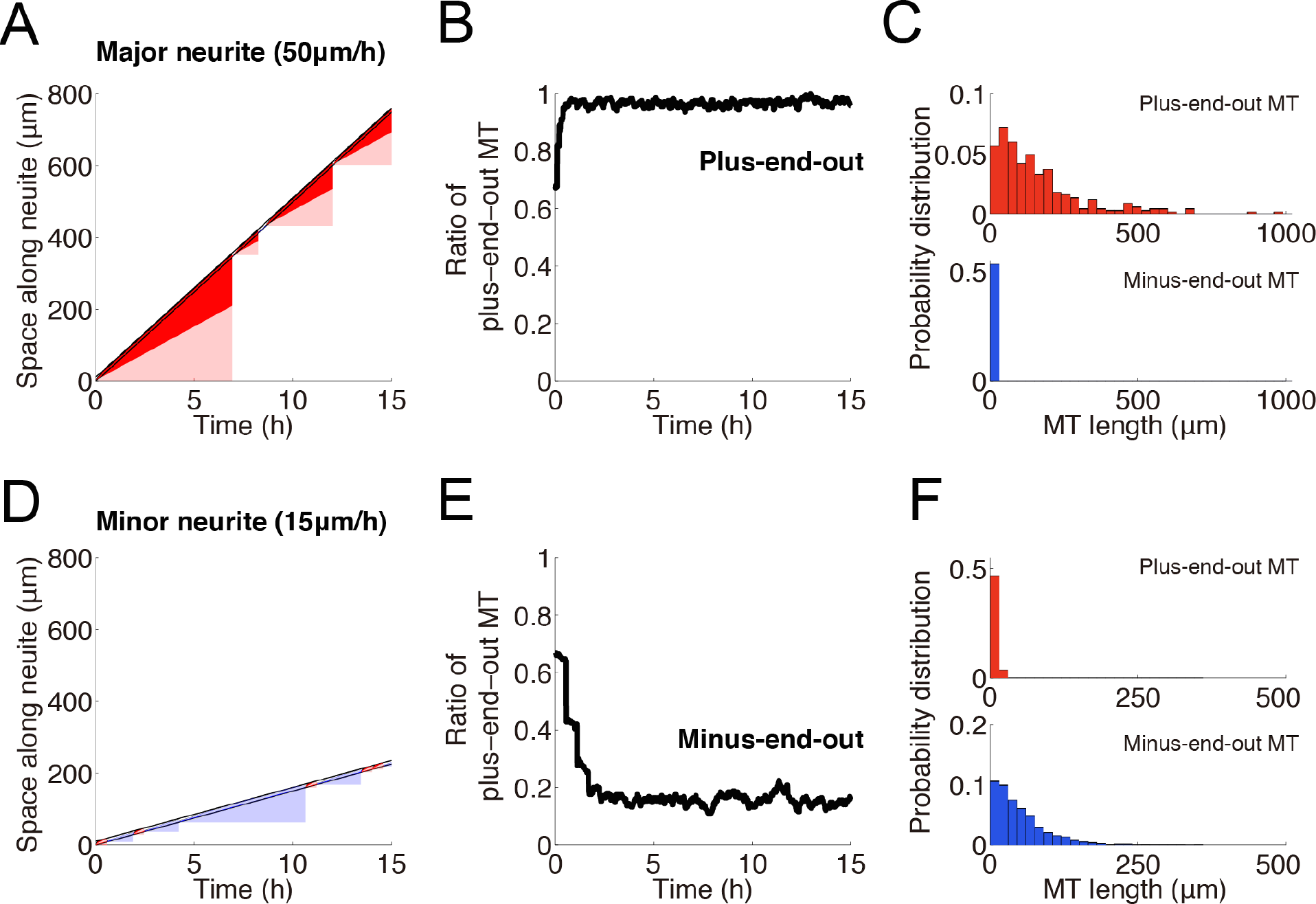
Self-organized MT orientations in invertebrate axon and dendrites. The temporal dynamics of MT replacements in fast- **(A-C)** and slow- **(D-F)** growing neurites were simulated with a high hydrolysis speed (30 (µm/h)). All panels correspond to the ones in Fig. 4.

### Unified view of three distinct MT orientations

Finally, we examined the phase diagram of the MT orientation patterns depending on the neurite growth speed and the hydrolysis speed (the speed of GTP cap disappearance) (**Fig. 6**). The plus-end-out pattern emerged when the neurite growth was faster than the growth of the minus end and slower than that of the plus end (**Fig. 6(++)**). When the neurite growth was slower than the GTP cap disappearance, all MTs in the growth cone were extinguished, which was biologically unrealistic. When the neurite growth was slower than that of the minus end, either the mixed or minus-end-out pattern was achieved depending on the balance between hydrolysis and neurite growth. We obtained the mixed pattern (**Fig. 6(−+)**) and the minus-end-out pattern (**Fig. 6(−−)**) when the neurite growth was faster and slower than the GTP cap disappearance, respectively. In summary, this phase diagram provides a unified view of three distinct MT orientation patterns.

**Figure 6:**
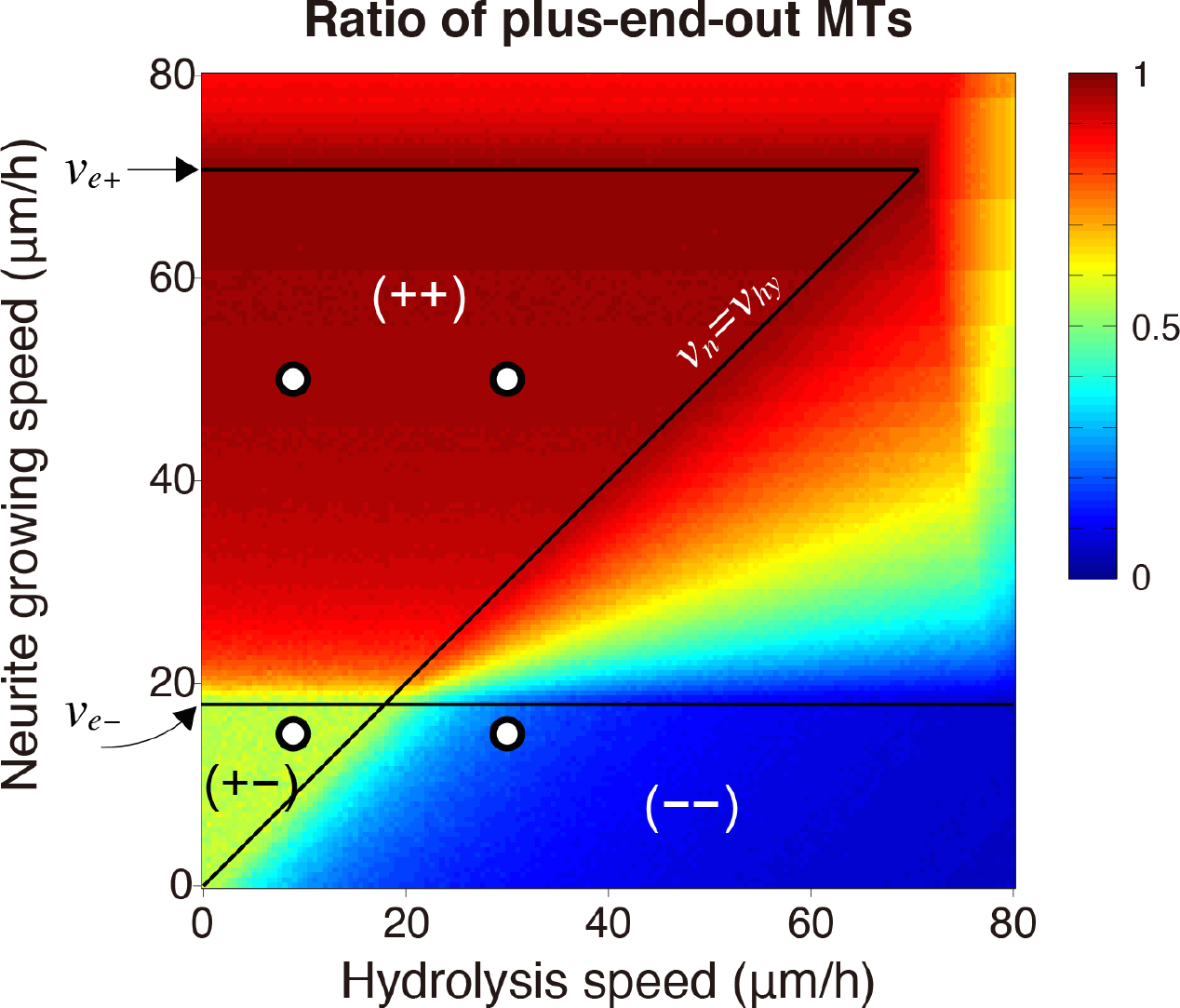
Phase diagram of MT orientation patterns. The MT orientation patterns depending on the neurite growth speed and hydrolysis speed were examined. The heat map indicates the proportion of the total length of plus-end-out MTs. (++), (−+) and (−−) represent regions of the plus-end-out, minus-end-out and mixed MT orientation patterns, respectively. Four open circles correspond to the plus-end-out and mixed patterns in Fig. 4 and the plus-end-out and minus-end-out patterns in Fig. 5.

## Discussion

We have presented a biophysical computational model of the MT kinetics in a growing neurite to reveal how neurites acquire distinct MT orientation patterns, which specify the identities of the axon and dendrites. The model was constructed by the elongation and catastrophic shrinkage of MTs in the growth cone and replacement of the existing MTs with actively transported MT fragments. The model successfully generated the plus-end-out, minus-end-out and mixed patterns of MT orientation observed in the axon and in vertebrate and invertebrate dendrites, respectively. We also determined how those three patterns emerged depending on the balance between neurite growth speed and the hydrolysis rate.

### Model validity

In our model, for the sake of simplicity, we adopted the following assumptions. First, we assumed that the neurite growth speed could be given as a control parameter independent of the MTs. In other words, MTs were passive and did not actively contribute to neurite protrusion, even though the MTs do interact with F-actin, which drive the growth cone motility [20]. At least, in the current state of our knowledge, it is intractable to model how the MTs regulate neurite protrusion via interaction with F-actin in the growth cone. Nevertheless, regardless of the interaction between MT and F-actin, our model demonstrated that the neurite growth speed was an essential factor to specify the MT orientation pattern. Second, we assumed MT replacement, in which inactivation of the existing MT and insertion of the new MT fragment into the MT bundle occur at the same time, leading to a fixed number of MTs. It can be speculated that those events must occur asynchronously in reality. In addition, the number of MTs has been reported to differ between major and minor neurites [29]. In this study, we avoided modeling the underlying processes due to a lack of knowledge. However, our model still had the power to predict the fractions of plus- and minus-end-out MTs, even if the number of MTs varies. Therefore, the minimalist model we developed had biological plausibility and was informative enough to provide a unified view of three types of MT orientations.

### Model predictions

On the basis of the model, we have here provided several experimentally testable predictions. According to the phase diagram (**Fig. 6**), whether neurites acquire a plus-end-out pattern, i.e., axon identity, was determined by the neurite growth speed relative to the elongation speeds of MT at the plus and minus ends, which depend on tubulin concentration. Thus, if tubulin concentration increases, the plus-end-out pattern could convert to the mixed or minus-end-out pattern because MTs at both the plus and minus ends would accelerate and catch up to the neurite growth. Conversely, a decrease in tubulin concentration could lead to conversion from the mixed and minus-end-out patterns to the plus-end-out pattern. Such conversions between the MT orientation patterns could also be induced by overexpressing CAMSAPs, which modulate the elongation speeds of the MT at both ends [30]. In addition, the phase diagram showed that the hydrolysis rate was key in determining whether minor neurites acquire the mixed or minus-end-out patterns. Thus, a change in the hydrolysis rate could induce conversion between vertebrate- and invertebrate-type dendrites. Moreover, our model predicted that the co-existence of minus-end-out and mixed patterns was impossible under the same MT kinetic parameters. Therefore, we have offered a biologically feasible model, but further experimental investigation is needed.

### Localizations of tau and MAP2

It is worth mentioning two MT-associated proteins (MAPs), which are well known as molecular markers for the axon and dendrites: tau is distributed throughout the neurons and enriched in the axon terminal, whereas MAP2 is specifically localized in the dendritic shaft [31]. As a mechanism for the polarized distributions, it can be speculated that tau and MAP2 bundle parallel and anti-parallel MTs, respectively, but this possibility has not been verified. In addition, the polarized distribution was thought not to be important for the acquisition of axon and dendrite identities, because it was obtained after the self-organization of MT orientations [7]. Moreover, studies on non-neural cells showed that the overexpression of the MAPs induced extension of the processes involving plus-end-out MTs, irrespective of tau or MAP2 [32,33]. A recent study, nevertheless, demonstrated that the local application of semaphorin 3A to the axonal growth cone induced the redistribution of MAPs, which then initiated the conversion from axon to dendrite. This finding suggested an important role of MAPs in the acquisition of axon and dendrite identities [34]. Thus, further investigation is needed to understand the whole picture of neuronal polarization, and we hope that our simple model will inspire future studies by other researchers.

## Materials and Methods

We mathematically described the biophysical model. The model neurite consisted of a neurite shaft and a growth cone with length *L_g_* at the neurite tip. The neurite growth was expressed by

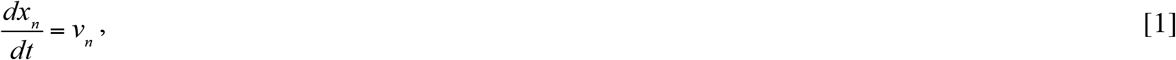

where *x_n_* and *v_n_* indicate the position and growth speed of the neurite tip, respectively. The elongation of the MT was described by

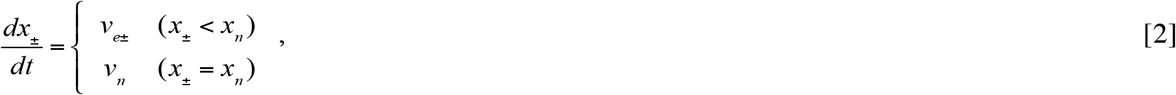

where *x_+_* and *x_−_* indicate the positions of the plus and minus end of the MTs pointing to the neurite terminal, respectively, and *v_e+_* and *v_e−_* indicate the elongation speed of the MTs at the plus and minus ends, respectively. Note that *v_e±_* was determined by

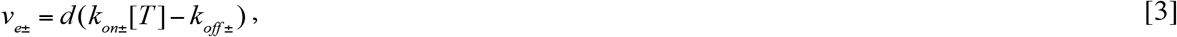

where [*T*] and *d* indicate the tubulin concentration in the growth cone and the length increment of single polymerization, respectively. We assumed that free tubulin was abundant and its concentration constant. In the model, when the MT reaches and contacts the neurite terminal (*x_±_* = *x_n_*), the MT accompanies the neurite growth. During catastrophe caused by the loss of the GTP cap at the plus end, the MT shrinks according to the equation

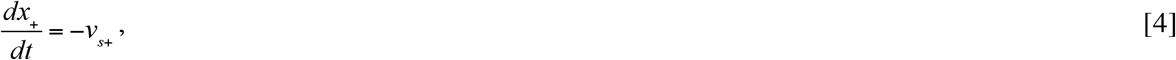

where *v_s+_* indicates the speed of the catastrophic shrinkage. In the model, we adopted vectorial hydrolysis [23], in which hydrolysis occurred only at the interface between the GTP- and GDP-bound tubulins. The hydrolysis was described by

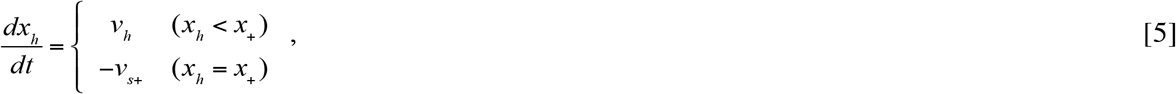

where *x_h_* and *v_h_* indicate the position and speed of the GTP/GDP interface, respectively. Note that *v_s+_* was determined by *v_s+_* = *dk_hy_*, where *k_hy_* indicates hydrolysis rate. Once the GTP cap at the plus end disappeared (*x_h_* = *x_+_*), the MT started catastrophic shrinkage. The MT then exited the growth cone (*x_+_* < *x_n_−L_g_*) and was replaced by a randomly oriented MT fragment. The elongating MTs were also spontaneously replaced by randomly oriented MT fragments in the growth cone at a rate of *k_rep_*. The parameters used here are listed in **Table. 1**.

**Table 1:** Parameter values published in the literature and used in the simulations.

| Parameter | Value | Reference |
| --- | --- | --- |
| $k_{on+}$ | 3.2 ($\mu\text{M}^{-1}\text{s}^{-1}$) | Howard et al. [35] |
| $k_{off+}$ | 0.1 ($\text{s}^{-1}$) | Howard et al. [35] |
| $k_{on-}$ | 0.83 ($\mu\text{M}^{-1}\text{s}^{-1}$) | – |
| $k_{off-}$ | 0.2 ($\text{s}^{-1}$) | – |
| $k_{cat}$ | 290 ($\text{s}^{-1}$) | Howard et al. [35] |
| $k_{hy}$ | 4 ($\text{s}^{-1}$) | Ranjith et al. [26] |
| $[T]$ | 10 ($\mu\text{M}$) | – |
| $d$ | 8/13 (nm) | Howard et al. [35] |
| $v_{e+}^*$ | 70.7 ($\mu\text{m}/\text{h}$) | Estimated by $d(k_{on+}[T] - k_{off+})$<br>Consistent with Hendershott and Vale [30] |
| $v_{e-}^*$ | 17.9 ($\mu\text{m}/\text{h}$) | Estimated by $d(k_{on-}[T] - k_{off-})$<br>Consistent with Hendershott and Vale [30] |
| $v_{s+}^*$ | 642.5($\mu\text{m}/\text{h}$) | Estimated by $dk_{cat}$ |
| $v_{hy}^*$ (vertebrate condition) | 8.86 ($\mu\text{m}/\text{h}$) | Estimated by $dk_{hy}$ |
| $v_{hy}^*$ (invertebrate condition) | 30 ($\mu\text{m}/\text{h}$) | – |
| $k_{rep}^*$ | 0.005 ($\text{min}^{-1}$) | – |
| $v^*$ (major neurite) | 50 ($\mu\text{m}/\text{h}$) | Toriyama et al. [13] |
| $v^*$ (minor neurite) | 15 ($\mu\text{m}/\text{h}$) | – |
| $L_g^*$ | 10 ($\mu\text{m}$) | – |

Parameters indicated by (\*) were used in simulations.

## Authors’ contributions

H.N. conceived the project and developed the models. H.N. and K.U. performed the computational simulation. H.N. wrote the manuscript, and S.I. proofread its final version.

## Acknowledgments

We are grateful to Dr. Michiyuki Matsuda for his valuable comments. We also thank Dr. Masataka Yamao for critically reviewing the manuscript.

## References

1. Arimura N, Kaibuchi K. 2007 Neuronal polarity: from extracellular signals to intracellular mechanisms. Nat. Rev. Neurosci. 8, 194–205. (doi:10.1038/nrn2056)

2. Naoki H, Nishiyama M, Togashi K, Igarashi Y, Hong K, Ishii S. 2016 Multi-phasic bi-directional chemotactic responses of the growth cone. Sci. Rep. 6, 36256. (doi:10.1038/srep36256)

3. Conde C, Cáceres A. 2009 Microtubule assembly, organization and dynamics in axons and dendrites. Nat. Rev. Neurosci. 10, 319–332. (doi:10.1038/nrn2631)

4. Kapitein LC, Hoogenraad CC. 2015 Building the Neuronal Microtubule Cytoskeleton. Neuron. 87, 492–506. (doi:10.1016/j.neuron.2015.05.046)

5. van Beuningen SFB, Hoogenraad CC. 2016 Neuronal polarity: Remodeling microtubule organization. Curr. Opin. Neurobiol. 39, 1–7. (doi:10.1016/j.conb.2016.02.003)

6. Kapitein LC, Hoogenraad CC. 2011 Which way to go? Cytoskeletal organization and polarized transport in neurons. Mol. Cell. Neurosci. 46, 9–20. (doi:10.1016/j.mcn.2010.08.015)

7. Baas PW, Lin S. 2011 Hooks and comets: The story of microtubule polarity orientation in the neuron. Dev. Neurobiol. 71, 403–418. (doi:10.1002/dneu.20818)

8. Kwan AC, Dombeck D a, Webb WW. 2008 Polarized microtubule arrays in apical dendrites and axons. Proc. Natl. Acad. Sci. U. S. A. 105, 11370–11375. (doi:10.1073/pnas.0805199105)

9. Kleele T et al. 2014 An assay to image neuronal microtubule dynamics in mice. Nat. Commun. 5, 4827. (doi:10.1038/ncomms5827)

10. Yau KW, Schatzle P, Tortosa E, Pages S, Holtmaat A, Kapitein LC, Hoogenraad CC. 2016 Dendrites In Vitro and In Vivo Contain Microtubules of Opposite Polarity and Axon Formation Correlates with Uniform Plus-End-Out Microtubule Orientation. J. Neurosci. 36, 1071–1085. (doi:10.1523/JNEUROSCI.2430-15.2016)

11. Stone MC, Roegiers F, Rolls MM. 2008 Microtubules have opposite orientation in axons and dendrites of Drosophila neurons. Mol. Biol. Cell 19, 4122–9. (doi:10.1091/mbc.E07-10-1079)

12. Goodwin PR, Sasaki JM, Juo P. 2012 Cyclin-Dependent Kinase 5 Regulates the Polarized Trafficking of Neuropeptide-Containing Dense-Core Vesicles in Caenorhabditis elegans Motor Neurons. J. Neurosci. 32, 8158–8172. (doi:10.1523/JNEUROSCI.0251-12.2012)

13. Toriyama M, Sakumura Y, Shimada T, Ishii S, Inagaki N. 2010 A diffusion-based neurite length-sensing mechanism involved in neuronal symmetry breaking. Mol. Syst. Biol. **6**. (doi:10.1038/msb.2010.51)

14. Samuels DC, Hentschel HG, Fine A. 1996 The origin of neuronal polarization: a model of axon formation. Philos. Trans. R. Soc. Lond. B. Biol. Sci. 351, 1147–1156. (doi:10.1098/rstb.1996.0099)

15. Fivaz M, Bandara S, Inoue T, Meyer T. 2008 Robust Neuronal Symmetry Breaking by Ras-Triggered Local Positive Feedback. Curr. Biol. 18, 44–50. (doi:10.1016/j.cub.2007.11.051)

16. Naoki H, Nakamuta S, Kaibuchi K, Ishii S. 2011 Flexible search for single-axon morphology during neuronal spontaneous polarization. PLoS One 6. (doi:10.1371/journal.pone.0019034)

17. Takano T et al. In press. Discovery of long-range inhibitory signaling to ensure single axon formation. (doi:10.1038/s41467-017-00044-2)

18. Nonaka S, Naoki H, Ishii S. 2011 A multiphysical model of cell migration integrating reaction-diffusion, membrane and cytoskeleton. Neural Networks 24, 979–989. (doi:10.1016/j.neunet.2011.06.009)

19. Dent EW, Gupton SL, Gertler FB. 2011 The growth cone cytoskeleton in Axon outgrowth and guidance. Cold Spring Harb. Perspect. Biol. 3, 1–39. (doi:10.1101/cshperspect.a001800)

20. Dehmelt L, Halpain S. 2004 Actin and Microtubules in Neurite Initiation: Are MAPs the Missing Link? J. Neurobiol. 58, 18–33. (doi:10.1002/neu.10284)

21. Mitchison T, Kirschner M. 1984 Microtubule assembly nucleated by isolated centrosomes. Nature 312, 232–7. (doi:10.1038/312232a0)

22. Akhmanova A, Hoogenraad CC. 2015 Microtubule minus-end-targeting proteins. Curr. Biol. 25, R162–R171. (doi:10.1016/j.cub.2014.12.027)

23. Bowne-Anderson H, Zanic M, Kauer M, Howard J. 2013 Microtubule dynamic instability: A new model with coupled GTP hydrolysis and multistep catastrophe. BioEssays 35, 452–461. (doi:10.1002/bies.201200131)

24. Hinow P, Rezania V, Tuszyński JA. 2009 Continuous model for microtubule dynamics with catastrophe, rescue, and nucleation processes. Phys. Rev. E - Stat. Nonlinear, Soft Matter Phys. **80**. (doi:10.1103/PhysRevE.80.031904)

25. Mazilu I, Zamora G, Gonzalez J. 2010 A stochastic model for microtubule length dynamics. Phys. A Stat. Mech. its Appl. 389, 419–427. (doi:10.1016/j.physa.2009.10.017)

26. Ranjith P, Lacoste D, Mallick K, Joanny JF. 2009 Nonequilibrium self-assembly of a filament coupled to ATP/GTP hydrolysis. Biophys. J. 96, 2146–2159. (doi:10.1016/j.bpj.2008.12.3920)

27. Vorobjev I a, Rodionov VI, Maly I V, Borisy GG. 1999 Contribution of plus and minus end pathways to microtubule turnover. J. Cell Sci. 112 (Pt 1), 2277–2289.

28. Schaefer AW, Kabir N, Forscher P. 2002 Filopodia and actin arcs guide the assembly and transport of two populations of microtubules with unique dynamic parameters in neuronal growth cones. J. Cell Biol. 158, 139–152. (doi:10.1083/jcb.200203038)

29. Seetapun D, Odde DJ. 2010 Cell-length-dependent microtubule accumulation during polarization. Curr. Biol. 20, 979–988. (doi:10.1016/j.cub.2010.04.040)

30. Hendershott MC, Vale RD. 2014 Regulation of microtubule minus-end dynamics by CAMSAPs and Patronin. Proc. Natl. Acad. Sci. 111, 5860–5865. (doi:10.1073/pnas.1404133111)

31. Kosik KS, Finch EA. 1987 MAP2 and tau segregate into dendritic and axonal domains after the elaboration of morphologically distinct neurites: an immunocytochemical study of cultured rat cerebrum. J. Neurosci. 7, 3142–53.

32. Chen J, Kanai Y, Cowan NJ, Hirokawa N. 1992 Projection domains of MAP2 and tau determine spacings between microtubules in dendrites and axons. Nature 360, 674–677. (doi:10.1038/360674a0)

33. Baas PW, Pienkowski TP, Kosik KS. 1991 Processes induced by tau expression in Sf9 cells have an axon-like microtubule organization. J. Cell Biol. 115, 1333–1344. (doi:10.1083/jcb.115.5.1333)

34. Nishiyama M, Togashi K, von Schimmelmann MJ, Lim C-S, Maeda S, Yamashita N, Goshima Y, Ishii S, Hong K. 2011 Semaphorin 3A induces CaV2.3 channel-dependent conversion of axons to dendrites. Nat. Cell Biol. 13, 677–686. (doi:10.1038/ncb2255)

35. Howard J. 2001 Mechanics of Motor Proteins and the Cytoskeleton. (doi:10.1017/CBO9781107415324.004)

